# CoBRA: Compound Binding Site Prediction using RNA Language Model

**DOI:** 10.1101/2025.09.03.673903

**Authors:** Wonkyeong Jang, Woong-Hee Shin

## Abstract

Ribonucleic acid (RNA) molecules perform a variety of functions within cells and thus implicated in various human diseases such as cancer. The fact that the proteins constitute a small portion of mRNAs and the ability of RNA to form highly specific three-dimensional binding pockets for small molecules have generated considerable interest in developing therapeutic agents that target RNAs as an alternative target for small-molecule drugs. Thus, like proteins, precise prediction of small-molecule binding sites across different classes of RNA targets could act as an starting point for drug discovery.

In this study, we present a lightweight deep learning framework called Compound Binding Site Prediction for Ribonucleic Acid (CoBRA). This framework can predict ligand-binding nucleotides without requiring any explicit structural information. Our approach uses residue-level embeddings obtained from a pre-trained RNA language model. These embeddings encapsulate the contextual and statistical properties of each nucleotide and are used in a frozen state as the input for a multi-layer perceptron classifier performing binary classification of binding versus non-binding residues. The model was trained using combinations of ten distinct RNA language models and six different loss functions, using the TR60 and HARIBOSS datasets, and tested on four independent benchmark sets (RB9, JL10, TL12, and TE18). The performance of CoBRA demonstrates a relative improvement of 22.1% in Matthew correlation coefficient and an increase in sensitivity of 45.6% compared to existing state-of-the-art RNA– ligand binding site prediction methods with structural information. These results show that sequence-based language model embeddings alone, which do not require any explicit coordinate or distance information, can match or outperform structure-based methods. This makes it a flexible tool for predicting binding sites across diverse RNA targets. CoBRA is available at https://github.com/WonkyeongJang/CoBRA.

## Introduction

RNA molecules play diverse roles in the cell’s gene regulation, translation, and structural organization. They are also involved in various human diseases, including cancer, neurological disorders, cardiovascular dysfunction, and developmental abnormalities [1–3]. Given that only 1.5% of mRNAs are translated into protein and 80% of proteins are predicted to be undruggable, the targeting of mRNAs as a therapeutic target has recently been highlighted. Additionally, there has been an escalating focus on their potential as a small-molecule drug targets, owing to their capacity to generate structured regions for binding with high specificity [4,5]. These RNA-ligand interactions present novel avenues for therapeutic intervention and thus a precise computational prediction of ligand-binding sites across diverse RNA categories are needed.

A variety of computational methods have been developed to predict RNA–ligand binding sites, and most of them rely on structural information either directly or indirectly. Rsite [6] calculates Euclidean distances between nucleotides based on RNA tertiary structures and predicts binding sites if the nucleotides are located at extreme points of the distance curve. Rsite2 [7] extends the strategy to secondary structures by computing Hamming distances, offering a coarse-grained manner incorporating structural information indirectly. Rbind [8] represents RNA tertiary structures as a network, where nucleotides are treated as nodes and non-covalent spatial interactions form edges. Binding site prediction is then performed based on network centrality metrics such as degree and closeness. RNAsite [9] is a machine learning-based method that extracts sequence and structure-derived features of nucleotides with a sliding window strategy. A random forest classifier is employed to predict binding status based on these features. RLBind [10] integrates both a full-length RNA sequence and structural features to predict binding site probabilities at the nucleotide level using a deep learning framework. Global features such as network topological properties, biochemical characteristics, and accessible surface areas, are processed by convolutional layers. On the other hand, local contextual features, including nucleotide types and evolutionary conservation, are captured using a dense neural network. ZHmolReSTasite (ZeSTa) [11] utilizes point clouds of solvent-excluded surfaces generated from RNA tertiary structures. These are then converted into normalized topographic images corresponding to individual nucleotides. The resulting representations are used as input features to a deep learning model that learns surface-based geometric features.

In this study, we propose a lightweight deep learning-based model, called CoBRA (<u>Co</u>mpound <u>B</u>inding Site Prediction for <u>R</u>N<u>A</u>), that predicts RNA–ligand binding sites using an RNA language model (LM), not relying on the RNA structural information. The model utilizes residue-level embeddings derived from pre-trained RNA language models, which implicitly encode the contextual and statistical properties of each nucleotide. These embeddings are used in a frozen state and subsequently entered into a multi-layer perceptron (MLP) classifier for residue-level binary classification. The proposed framework is designed to operate without explicit structural features such as 3D coordinates or distance metrics, and its modularity allows for flexible experimentation. A combination of ten RNA LMs and six loss functions was systematically trained, combining TR60 [9] and HARIBOSS set [12], and then evaluated to identify optimal configurations for the RNA-ligand binding site prediction. The final model was evaluated in four benchmark sets: TE18 [9], RB9, JL10, and TL12 [11]. CoBRA achieved a relative improvement of approximately 22.1% in MCC and 45.6% in recall compared to the state-of-the-art RNA-ligand binding site prediction programs.

## Method

### Dataset Preparation

The prediction of RNA–ligand binding sites is a binary classification task at the residue level. The model predicts the binding of each nucleotide (residue) in a given RNA sequence to an organic small molecule or metal ion. Each residue is designated as binding (1) or non-binding (0). The labeling of a nucleotide is labeled as a binding site residue in a given co-crystalized structure of an RNA and a compound contingent upon the minimum Euclidean distance between any of its atoms and any ligand atom being less than or equal to 4 Å. However, if the interacting molecule is one of non-specific experimental additives such as water, SO_4_^2-^, and PO_4_^3-^, then the complex structure is excluded from the datasets.

This study utilized a total of six publicly available RNA–ligand complex structure datasets, including HARIBOSS, TR60, RB9, TL12, JL10, and TE18. The construction of the HARIOBSS set was achieved by Panei et al. [12]. This process entailed the extraction of RNA-small molecule complexes from the PDB. Subsequently, these complexes were clustered based on their sequence and structural similarity with RNA. Su et al. [9] also collected 712 RNA-small molecule crystal structures and used TM-scoreRNA to calculate their structural similarity, resulting in 72 representatives. The dataset is further segmented into two distinct sets: the TR60 set and the TE18 set. These sets are utilized for the training and testing of RNAsite. Gao et al. generated RB9, TL12, and JL10 datasets to assess their binding site prediction program, ZeSTa [11]. RB19 was originally designed to test the performance of RBind [8] and RNAsite [9]. The authors removed ten structures from the dataset that overlapped with TR60 and renamed the set as RB9. From the RNA-small molecule structures deposited after January 2021, JL10 is characterized by junction loop structures, which exhibit high structural complexity. In contrast, TL12 structures are devoid of the loop, resulting in low structural complexity. Among the six datasets, HARIBOSS and TR60 were combined for training and validation, yielding 432 structures. The dataset was segmented using a pre-determined random seed to create three subsets with a ratio of 8:1:1 for training, validation, and internal testing, respectively. The remaining four, RB9, JL10, TL12, and TE18, were reserved as test sets. To ensure independent evaluation, any overlapping sequences between the training and test sets were removed. The problem formulation and dataset design enable quantitative assessment of both prediction accuracy and generalizability under practically meaningful scenarios.

### Model Architecture

To predict whether each nucleotide in an RNA sequence is a binding site or not, a residue-level binary classification model was designed. Figure 1 illustrates the model architecture of CoBRA. The program’s input is a query RNA sequence which is embedded using pre-trained RNA LMs. During the training process, these embeddings are maintained in a frozen state and do not undergo updates.

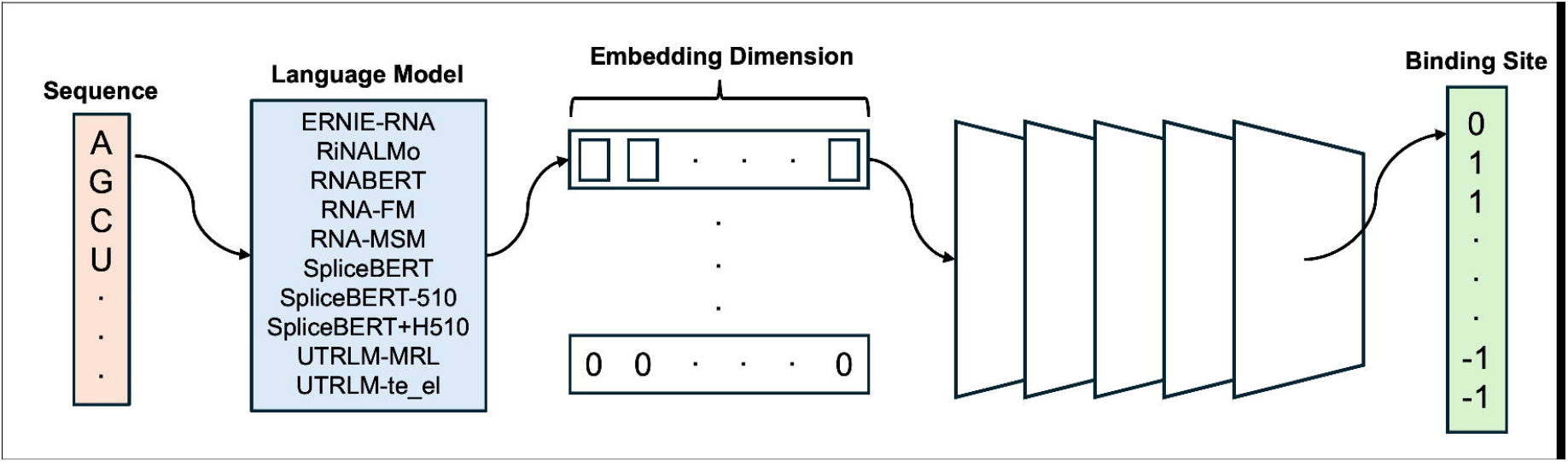

The prediction model employs a multi-layer perceptron (MLP) architecture comprising five fully connected layers (see Figure 1). Each layer is followed sequentially by layer normalization, a ReLU activation function, and dropout with a probability of 0.1, promoting stable training and improved generalization. The final output layer produces a two-dimensional logit for each residue, corresponding to the binding and non-binding classes. These are converted into probabilities via a softmax function.

All inputs are standardized to a maximum sequence length of 161 residues. Sequences of shorter length are padded with zeros in the input matrix X and with -1 in the target vector y. A masking strategy is implemented to ensure that the padding regions do not contribute to the loss computation or affect model training.

The model training was executed using the PyTorch [13] framework. The optimization process was executed employing the AdamW [14] optimizer, with an initial learning rate set to 5 × 10^-4^. The cosine annealing method was employed for the purpose of learning rate scheduling. All experiments were run with a batch size of 4, a dropout probability of 0.1, for 100 epochs, and with the random seed fixed to 42 to ensure reproducibility.

### RNA Language Models

Ten pre-trained RNA LMs were employed to pick up the best model for residue-level binding site prediction. The models were pre-trained on diverse RNA types for various learning objectives, resulting in differences in embedding dimensionality and representational properties. Table 1 lists the RNA LMs, their pre-training targets, and embedding size.

The models can be categorized into two groups based on pre-training sets. The first category is the models trained on non-coding RNAs (ncRNAs), composed of ERNIE-RNA [15], RiNALMo [16], RNABERT [17], RNA-FM [18], and RNA-MSM [19]. This group captures generalizable structural patterns across various ncRNA families. ERNIE-RNA [15], comprised of 12 Transformer blocks, is pre-trained via masked language modeling (MLM) on 20 million ncRNAs from RNAcentral with structural information. RiNALMo [16] is also pre-trained on 36 million ncRNAs, employing MLM with 33 Transformer blocks. RNABERT [17] adopts the pre-training BERT algorithm to 762 K ncRNAs. It also encodes the characteristics of the RNA family and structure. RNA-FM [18] is built upon 12 bidirectional Transformer encoder blocks that produce an L × 640 embedding matrix for input length L, pre-trained on 23.7 million ncRNAs. RNA-MSM [19] follows the MSA Transformer architecture with ten blocks to learn two-dimensional co-evolutionary signals from homologous multiple sequence alignments of 3,932 Rfam families.

The members of the second class are pre-trained on pre-mRNA or mRNA untranslated regions. SpliceBERT, SpliceBERT-510, SpliceBERT-H510 [20], UTRLM-MRL, and UTRLM-TE_EL [21] are members of the class. The category specializes in modeling sequence features in post-transcriptional regulatory regions. SpliceBERT [21] is a BERT-based model with six Transformer encoder layers pre-trained on 2 million RNA sequences from 72 vertebrate species. Additionally, two variants of the model were also employed: SpliceBERT-510, an intermediate checkpoint, and SpliceBERT-H510, pre-trained exclusively on human data. UTR-LM [21] is a six-layer Transformer encoder pre-trained via semi-supervised masked nucleotide reconstruction, 5′ UTR secondary-structure prediction, and minimum-free-energy regression on 5′ UTRs from multiple species. Two task-specific variants UTR-LM–TE_EL (translation efficiency and mRNA expression-level prediction) and UTR-LM–MRL (mean ribosome loading prediction) were also used to generate embeddings.

All language models were used in their pre-trained form with frozen parameters; no fine-tuning or weight updates were performed during training. Each RNA sequence was transformed into a residue-level embedding sequence using the corresponding model then padded to a fixed length of 161 residues before being fed into the MLP classifier. Embedding dimensionality varied across models, ranging from 120 to 1280. We systematically compared the predictive performance across different embedding types to assess their impact on RNA– ligand binding site prediction.

### Benchmarked Loss Functions

Compared to the RNA sequence length, the proportion of binding nucleotides (positive class) is low, leading to a severe class imbalance problem. To resolve the issue and quantitatively evaluate how different training objectives affect performance, we employed six loss functions.

Each loss function represents a different optimization strategy. Binary Cross-Entropy (BCE) serves as the conventional baseline for binary classification, encouraging predicted probabilities to converge to the ground truth. Class-balanced focal loss [22] and Tversky loss [23] are designed to address the class imbalance. The class-balanced focal loss emphasizes hard-to-classify samples by assigning them higher weights, while Tversky loss applies asymmetric weighting to false positives and false negatives, enabling recall-oriented optimization.

Dice loss [24] and Lovász hinge loss [25] focus on residue-level spatial alignment and structural consistency. Dice loss maximizes the overlap between predicted and true positive regions, whereas Lovász hinge loss optimizes the Intersection over Union directly.

Lastly, we introduce a composite loss that combines Triplet Center Loss (TCL) [26] and class-balanced focal loss to simultaneously improve class separation in the embedding space and address the class imbalance. This design is inspired by CLAPE-SMB [27], which demonstrated strong performance in protein-small molecule binding site prediction using a combination of TCL and class-balanced focal loss to enhance both feature discrimination and class imbalance handling. The total loss is defined as shown in Equation 1.

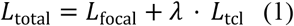

*L*_*focal*_ and *L*_tcl_ are class-balanced focal loss and TCL loss, respectively. λ is used as a weight to balance between the losses.

TCL learns a center vector for each class and encourages embeddings of the same class to cluster together while enforcing a margin-based separation between different classes (Equation 2).

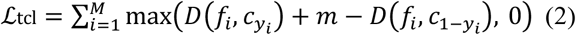

Here, *f* is the embedding vector of the input, *c*_*y*_ and *c*_*1−y*_ denote the center vectors of the true and opposite classes, respectively, and m is the margin. In our experiments, we set λ=0.2 and m=4. All combinations of the RNA LMs and loss functions were tried to make the prediction models and evaluated.

### Evaluation Metrics

To validate the models’ performance, we employed precision, recall, Matthews correlation coefficient (MCC), and the areas under the receiver operating characteristic and precision-recall curves (AUROC and AUPRC). To calculate the metric, the nucleotides were classified into four categories: true positive (TP), false positive (FP), true negative (TN), and false negative (FN). Positive and negative indicate whether a nucleic acid is predicted as a binding site, and true and false denote that the prediction is correct.

Precision and recall are the fraction of correct positive predictions among all positive calls and all actual positives, respectively (Equation 3,4).

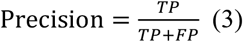

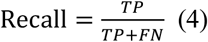

The MCC measures the quality of binary classifications even under class imbalance (Equation 5).

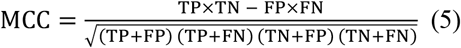

AUROC quantifies the model’s discrimination ability by measuring the area under the ROC curve (TP rate vs. FP rate), and AUPRC measures the area under the precision–recall curve, summarizing the trade-off between precision and recall across all thresholds.

### Laplacian-Based Curvature for RNA 3D Structure Analysis

To analyze and characterize the three-dimensional RNA structures quantitatively, Laplacian-based curvature descriptor (LN) [11] is employed. This metric quantifies the curvature at each nucleotide by measuring the deviation of its C3′ atom coordinates from the relative positions of surrounding residues. It thereby captures local structural distortion and reflects how naturally a nucleotide is embedded within the overall structure.

For each nucleotide, a coordinate vector was defined by C3′ atom position. Then a Gaussian kernel-based weighting function was constructed from the all residue pairwise Euclidean distance matrix following Equation 6.

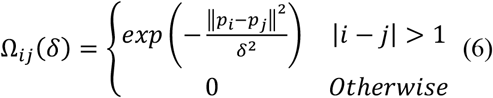

Here, δ is a single-scale parameter, set as the median of all pairwise distances between nucleotides in the structure. *p*_*i*_ and *p*_*j*_ denote the Euclidean coordinates of the C3′ atoms of nucleotides *i* and *j*. This choice of scale balances local and global shape sensitivity, avoiding excessive localization or over-smoothing.

Using the weight function, the Laplacian norm value *LN*_*i*_ for nucleotide *i* is defined as Equation 7.

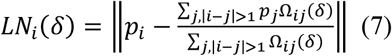

This value reflects how far nucleotide *i* lies from its Gaussian-weighted center, capturing the relative positional characteristics within the RNA structure. Higher LN values indicate convex or protruding regions, while lower values suggest concave or densely packed areas. The computed LN values were utilized for analysis and structural visualization.

## Result and Discussion

### Overall Performance Comparison and Model Selection

A total of 60 models were evaluated and compared, incorporating ten RNA LM embeddings and six loss functions. Table 2 lists the top ten MCC models benchmarked on the test set. A detailed data of individual models is given in Table S1.

The combination model that achieves the highest MCC is ERNIE-RNA with TCL focal as the language model and loss function, respectively. The model has also been determined to be second in terms of AUPRC and precision among the 60 models considered. A total of six RNA LMs were assessed in the study, and it was observed that ERNIE-RNA, RNA-FM, and RiNALMo emerged as top contenders, appearing in the top ten MCC models on three separate occasions. The three LMs were pre-trained from ncRNAs. Conversely, the LMs pre-trained on mRNA are not included among the top 10, with the exception of SpliceBERT-510. The comparatively diminished performance of mRNA-pretrained models might be due to their constrained exposure to a variety of RNA structures, which could result in diminished generalizability in binding site prediction when compared to ncRNA-pretrained counterparts.

A survey of the top ten MCC models reveals that nine of them utilize entropy-like loss functions, including focal, TCL focal, and BCE. Except for the three aforementioned functions, the Lovasz hinge with RiNALMo is ranked sixth in terms of MCC.

The distribution of MCC, AUPRC, and AUROC for all 60 models is illustrated in Figure 2 across the LMs and loss functions. As observed by the top ten MCC models, RiNALMo, ERNIE-RNA, and RNA-FM, which were pre-trained on ncRNAs, exhibited the highest average MCC, AUROC, AUPRC, and precision values (Figure 2A). With regard to loss functions, models that have been trained with BCE and focal loss have demonstrated a consistent and superior performance across key evaluation metrics in comparison to other loss functions. Conversely, models employing dice loss and Lovasz hinge loss exhibited unstable convergence and suboptimal performance in most cases (Figure 2B). In light of the collective performance, a selection of five models was made, and subsequent benchmarking was conducted on the external validation sets.

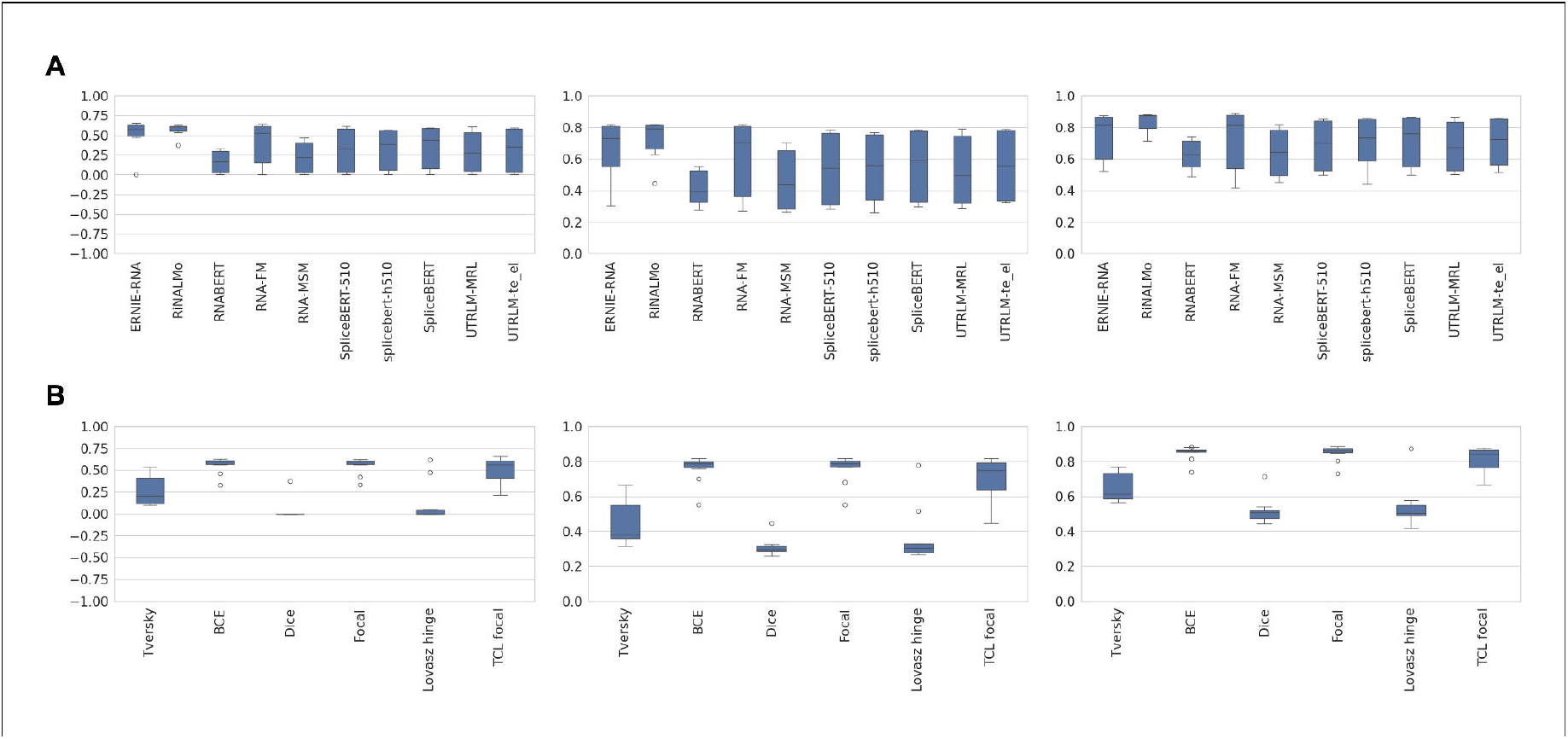

### Comparison with Other Existing Methods on Benchmark Sets

We conducted a benchmarking study of the top five MCC models on various test sets composed of RNA-compound crystal structures. RB9, JL10, TL12, and TE18. Table 3 presents the performance of the models and other RNA-compound binding site prediction methods. The CoBRA models are designated CoBRA-M1, M2, and so forth, following their MCC ranks.

While the CoBRA-M1, ERNIE-RNA combined with TCL focal model demonstrated the highest MCC, AUPRC, and precision among the five models evaluated on the test set, its performance was not optimal on the four benchmark sets. Conversely, CoBRA-M3, utilizing RiNALMo as the RNA LM and BCE as the loss function, exhibited the highest performance in MCC and AUROC on the average of the four test sets. For precision and recall, M1 and M2 performed best, respectively.

With regard to the MCC from individual benchmark sets, M2 has the highest value for JL10 and RB9, whereas M1 shows the best MCC for TL12. While M3 is only ranked top for TE18 among the five CoBRA models, it is ranked second for JL10, a set with a high structural complexity, and TL12, a set with a low structural complexity. This suggests that M3 has greater generalizability for RNA-compound binding site prediction problems than other models. Consequently, M3 is selected as a representative model for CoBRA, and the results are subjected to further analysis.

A comparative analysis of RNA-compound binding site prediction methods revealed that CoBRA exhibited superior performance, with the exception of TE18. On average, CoBRA exhibited a 15.4%, 9.2%, and 33.8% improvement in MCC, AUROC, and recall, respectively, when compared to ZeSTa, a state-of-the-art program. In particular, on datasets with relatively simpler structures, such as RB9 and TL12, CoBRA also outperformed other methods, with the exception of RB9 precision. In a similar vein, the CoBRA algorithm demonstrated the most optimal performance in all metrics for the highly complex structure set, JL10. Conversely, the TE18 dataset exhibited a decline in performance, with an MCC reduction of 41.90% in comparison to ZeSTa.

One of the successful cases of CoBRA is illustrated in Figure 3A. A microRNA of 125 bases in length forms a complex structure with bis-(3’,5’)-cyclic-dimeric-adenosine-monophosphate (PDB ID: 6WTR). A comparison of the two programs revealed that CoBRA exhibited superior performance in comparison with ZeSTa. The MCC, precision, and recall of CoBRA are 0.923, 0.946, and 0.946, respectively, while those of ZeSTa are 0.121, 0.455, and 0.142, respectively.

**Figure 3.**
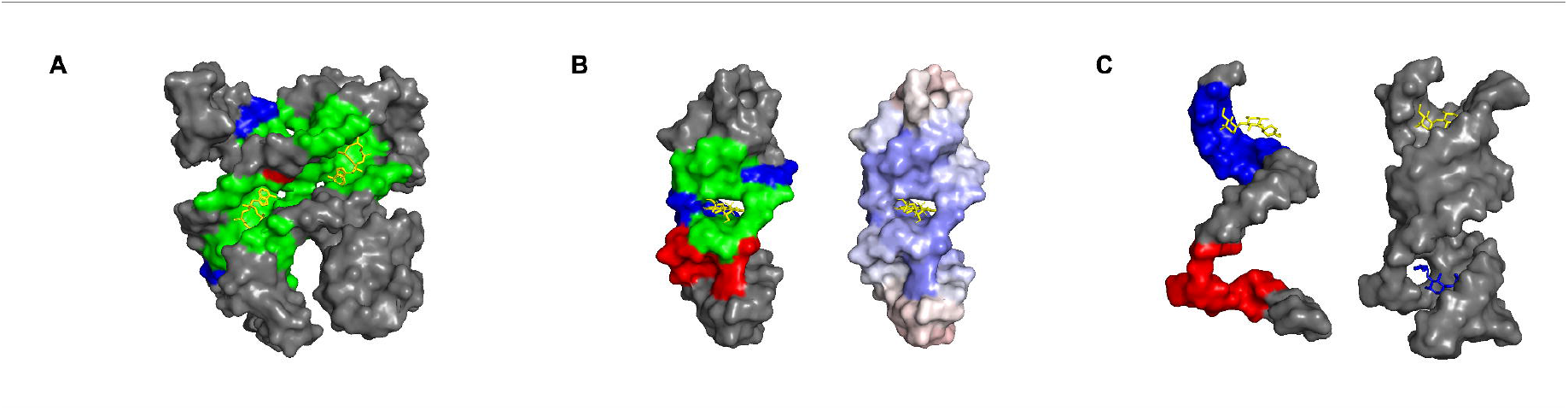
Case studies of CoBRA. Green, red, grey, and blue colors represent true positives, false positives, true negatives, and false negatives of CoBRA. **A**. A successfully predicted case of CoBRA. A pir-miRNA-300 complexed with bis-(3’,5’)-cyclic-dimeric-Adenosine-monophosphate (PDB ID: 6WTR). **B**. RNA Aptamer complexed with flavin mononucleotide (PDB ID: 1FMN). The right panel displays the distribution of Laplacian norm values, where blue indicates lower curvature (concave) and red indicates higher curvature (convex). **C**. Wrongly predicted case of CoBRA: sisomicin bound to bacterial ribosomal decoding site (PDB ID: 4F8U). The right panel shows an RNA homodimer, highlighting the inter-chain binding interface.

### Structure Analysis on TE18 Dataset

An investigation was conducted to examine the outcomes of TE18, wherein CoBRA exhibited the least optimal performance among the test sets. Dissecting the spatial structure of RNA (DSSR) [28] was employed to analyze the secondary structures of RNA crystal structures. The RNA secondary structures were classified into eight categories: stem, canonical, isolated canonical, internal loop, hairpin loop, burge, junction, and helix. A subsequent analysis of the prediction performance of CoBRA was conducted, with the structural type serving as the primary variable.

The program demonstrated a consistent level of accuracy, with a minimum of 51% for identifying binding sites across structural types. The highest accuracy was observed in junction sites, with an accuracy of 92%. On the other hand, the performance for identifying non-binding nucleotides exhibited variability according to structure, with higher accuracy in the stem (68%) and junction (67%), but lower accuracy in internal loops (47%) (Figure S1).

We also conducted a residue-level curvature analysis of the TE18 dataset by calculating the LN. As the LN values increase, the geometry of the residue evolves from concave to convex. The LN values of the binding and non-binding sites show distinct distribution (Figure S2), suggesting structural differences between the two classes. The binding site nucleotides have LN values ranging from 1.5 to 20.7, 10.9 on average. Conversely, the non-binding ones exhibit larger values, ranging from 2.3 to 23.0, with an average of 12.7. It can be inferred that the geometry of RNA compound binding sites is relatively concave, a finding that is also reported by Su et al. [9].

Despite the absence of explicit incorporation of structural characteristics such as concavity as an input feature in CoBRA, a statistically significant relationship was observed between the model predictions and the LN value, as indicated by a point-biserial correlation of -0.171 with a p-value of 2.8e-5. This demonstrates that a model trained solely on RNA sequence embeddings could explain structural characteristics via inherent sequence patterns. We also observed a correlation of -0.247 between LN value and the actual binding label, indicating that actual binding sites tend to be located in structurally concave regions. Correlation analyses with TP/TN/FP/FN from CoBRA prediction demonstrate that TP and TN exhibit relatively high correlation coefficients (TP: r = -0.188, p = 4.0e-6; TN: r = 0.260, p = 1.2e-10). No significant relationship was found for FP (r = -0.027, p = 0.505), and FN exhibited a modest negative correlation (r = -0.126, p = 2.0e-3), indicating a slight reduction in missed positive predictions. A notable example is provided in Figure 3B (PDB ID: 1FMN), illustrating the substantial correlation between CoBRA prediction and LN values. The flavin binding site of an RNA aptamer, defined by 4 Å from the co-crystalized ligand, exhibited a -0.541 correlation coefficient (p = 8.0e-4). The right panel displays the LN values of nucleotides, indicated by the color change from blue (concave) to red (convex). CoBRA demonstrates an accuracy in predicting the binding site, with a precision of 0.636. However, the program failed to accurately predict three nucleotides with low LN values (average: 8.36) and erroneously predicted four nucleotides with high LN values (average: 9.89). These results suggest that incorporating structural properties, such as secondary structure and curvature, as an additional input feature in the model input could yield an improved performance.

One of the most severe predicted cases of CoBRA in the TE18 dataset is sisomicin complexed with the bacterial ribosomal decoding site (PDB ID: 4F8U, Figure 3C). The crystal structure contains homodimer RNA chains, and two sisomicins bind to the interface of the dimer. For prediction, the RNA sequence of a single chain, B chain of the PDB structure, was used as the input. CoBRA demonstrated a successful prediction for one of the binding sites (the red-colored region of the left panel). However, given that the binding site is designated by the ligand with the same chain ID of the input RNA sequence (the blue region), the resulting accuracy for that particular sample was found to be 0.0. A visual inspection of both the A and B chains together (right panel) revealed that the region was misclassified as a false positive. This case demonstrates that the accuracy of prediction may be diminished when binding sites are situated at the interface with other chains of a query sequence. This phenomenon has also been observed in protein-ligand binding site prediction problems using LMs [29]. One potential solution to this issue involves the inference of structural information or the incorporation of multi-chain as an input.

### Prediction Results on Metal Binding Sites

Metal ions such as Mg^2+^ and Na^+^ often bind diffusely across the RNA surface to neutralize the negatively charged phosphate backbone, without forming specific binding pockets. These ions contribute to RNA structural stability and folding, as previously noted by Draper et al. [30]. In contrast, organic molecules typically bind within well-defined pockets, making their spatial localization and binding site prediction more tractable. Thus, predicting metal binding sites might be harder than predicting organic molecule binding sites because of the nature of the binding. Also, due to the electrostatic and steric nature of metal ion binding, they may associate with multiple structurally similar regions with comparable affinity, complicating the prediction task, which is reported in MetalionRNA [31].

The binding sites of the test sets were classified according to the type of binding molecule: metal ion or non-metallic compound. The accuracies of metal binding site prediction were 0.647, 0.531, 0.497, and 0.202 for RB9, JL10, TL12, and TE18, respectively. As anticipated, the non-metallic compound category demonstrated higher levels of accuracy: The values obtained for RB9, JL10, TL12, and TE18 were 0.733, 0.739, 0.690, and 0.468, respectively. This discrepancy is likely attributable to the inherent nature of RNA–metal ion interactions. The incorporation of physico-chemical characteristics has the potential to enhance the efficacy of predicting metal binding sites.

## Conclusion

Given the recent emphasis on RNA as a promising therapeutic target, the prediction of compound binding sites could serve as a fundamental starting point. The RNA binding site prediction models can be broadly categorized into two distinct approaches: structure-based and sequence-based. The structure-based approach leverages three-dimensional or secondary structural information to capture critical spatial and geometric features, while the sequence-based approach utilizes evolutionary and contextual patterns through sequence embeddings. While structure-based models generally demonstrate robust performance when accurate structural information is available, they exhibit significant performance decrement on complex RNA architectures, such as junction loops. Conversely, the sequence-based model proposed employs a more generalizable strategy.

The integration of ERNIE-RNA as an RNA LM and focal loss function has led to the development of a novel RNA-compound binding site program, called CoBRA. On various benchmark sets, the model demonstrated performance that was either superior to or comparable to contemporary state-of-the-art prediction methods. The program demonstrated its capacity for generalization across a range of external datasets, including RB9 and TL12, without requiring explicit structural inputs. This finding highlights the efficacy of sequence embeddings in capturing functional signals across a diverse array of RNA architectures.

Despite these advances, several limitations were identified through structural analysis on the TE18 set, which CoBRA exhibited suboptimal performance. The performance of the program is contingent upon the RNA secondary structure, denoted by DSSR. Also, it misses to predict binding site nucleotides located in concave regions, as indicated by low LN values. In addition, binding sites located at interfaces of RNA dimers were often missed when only a single RNA chain was provided as input. Looking forward, integrating lightweight structural features, such as LN, solvent accessible surface area, or secondary structure, with sequence-based embeddings holds promise for overcoming current limitations. Taken together, our findings establish a strong foundation for future research, suggesting that hybrid models leveraging both sequence and minimal structural information could achieve an improved performance in RNA-ligand binding site prediction.

