## Supplementary material for "CoBRA: Compound Binding Site Prediction using RNA Language Model": Figure S1

**Figure S1.** Confusion matrices of CoBRA depending on RNA secondary structures.

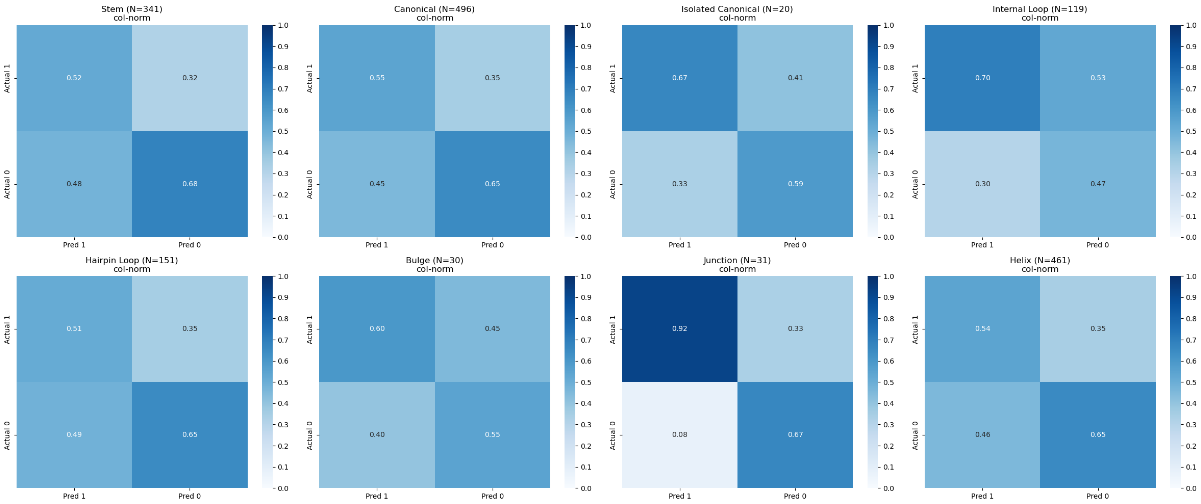

**Figure S2.** The distribution of Laplacian norm values of binding site (orange) and non-binding site (blue) nucleotides.

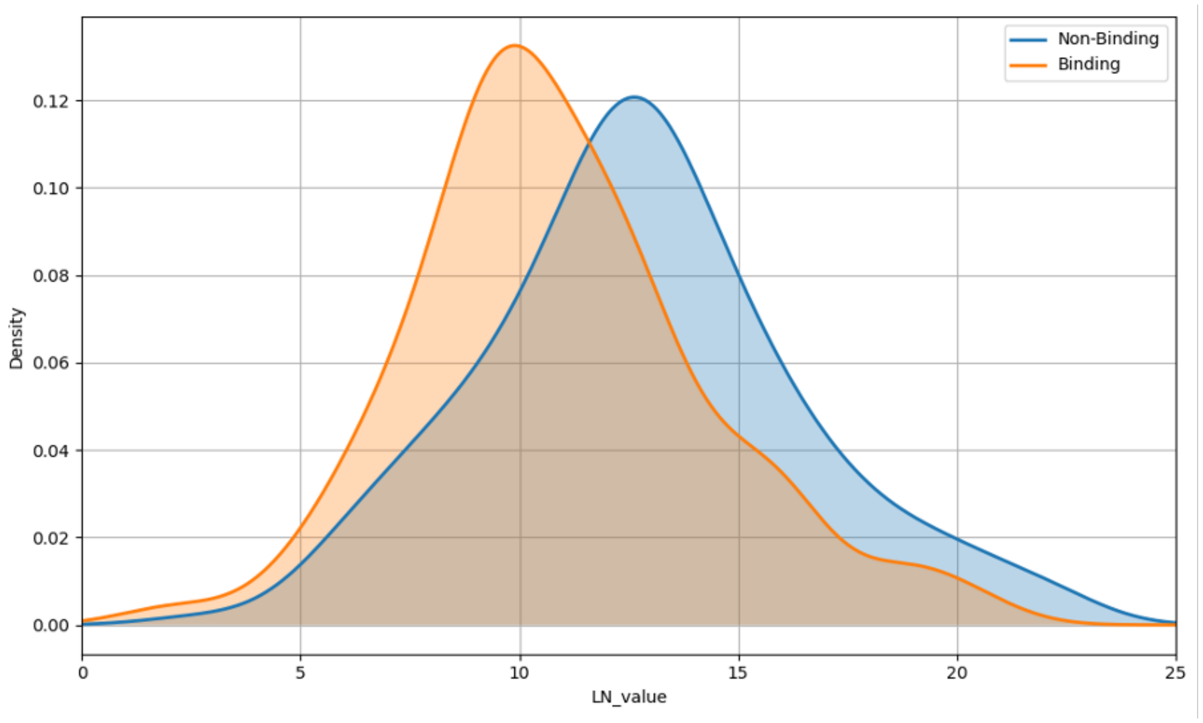
